# An Inter-Species Translation Model Implicates Integrin Signaling in Infliximab-Resistant Colonic Crohn’s Disease

**DOI:** 10.1101/776666

**Authors:** Douglas. K. Brubaker, Manu. P. Kumar, Paige. N. Vega, Austin. N. Southard-Smith, Alan. J. Simmons, Elizabeth. A. Scoville, Lori. A. Coburn, Keith. T. Wilson, Ken. S. Lau, Douglas. A. Lauffenburger

## Abstract

Anti-TNF therapy resistance is a major clinical challenge in Crohn’s Disease (CD), partly due to insufficient understanding of disease-site, protein-level mechanisms of CD and anti-TNF treatment resistance. Although some proteomics data from CD mouse models exists, data type and phenotype discrepancies contribute to confounding attempts to translate between preclinical animal models of disease and human clinical cohorts. To meet this important challenge, we develop and demonstrate here an approach called Translatable Components Regression (TransComp-R) to overcome inter-species and trans-omic discrepancies between CD mouse models and human subjects. TransComp-R combines CD mouse model proteomic data with patient pre-treatment transcriptomic data to identify molecular features discernable in the mouse data predictive of patient response to anti-TNF therapy. Interrogating the TransComp-R models predominantly revealed upregulated integrin pathway signaling via collagen-binding integrin ITGA1 in anti-TNF resistant colonic CD (cCD) patients. Toward validation, we performed single-cell RNA sequencing on biopsies from a cCD patient and analyzed publicly available immune cell proteomics data to characterize the immune and intestinal cell types contributing to anti-TNF resistance. We found that ITGA1 is indeed expressed in colonic T-cell populations and that interactions between collagen-binding integrins on T-cells and colonic cell types expressing secreted collagens are associated with anti-TNF therapy resistance. Biologically, TransComp-R linked previously disparate observations about collagen and ITGA1 signaling to a potential therapeutic avenue for overcoming anti-TNF therapy resistance in cCD. Methodologically, TransComp-R provides a flexible, generalizable framework for addressing inter-species, inter-omic, and inter-phenotypic discrepancies between animal models and patients to deliver translationally relevant biological insights.

**One Sentence Summary:** Brubaker *et al.* implicate dysregulated collagen-binding integrin signaling in resistance to anti-TNF therapy in Crohn’s Disease by developing a mouse-proteomic to human-transcriptomic translation model and confirm the associated inter-cellular signaling network using single-cell RNA sequencing.

## Introduction

Crohn’s disease (CD) is a chronic inflammatory bowel disease that may manifest in any region of the digestive tract. Anti-TNF therapeutics, including Infliximab (IFX) and Adalimumab (ADA), have emerged as remission-inducing therapies effective in 30-50% of patients, with up to 30% of these patients eventually developing secondary non-response (1). Because of this high rate of primary and secondary non-response to anti-TNF therapy, several studies have examined the signaling determinants of resistance through transcriptomic studies (1–7). However, transcriptomic characterization of anti-TNF resistance has yet to translate into effective strategies for overcoming therapeutic resistance, potentially due to the lack of functional proteomic characterization of Infliximab resistant CD prior to treatment. While some mouse studies have measured thousands of proteins by mass spectrometry and provided a detailed view of proteomic signaling in inflamed and uninflamed conditions (8, 9), these included no therapeutic stimuli making it challenging to generalize these therapy-independent murine signaling characterizations to clinical therapeutic resistance.

Translational Systems Biology aims to apply computational modeling to better translate biological insights from *in-vitro* and non-human *in-vivo* experimental models to the human disease context. Several recent studies have applied statistical and machine learning models to infer human disease-associated biology from model systems (10–13). Though these studies aimed at generalizable methods of interspecies translation, a limitation of these methods was the need for comparable molecular data types and similar phenotypes between model systems and humans. Therefore, these methods are not appropriate for translating the CD mouse model proteomic characterizations to understand Infliximab resistance in CD patients. If the challenges of interspecies, inter-omic translation between mismatched mouse and human phenotypes could be overcome, then the available mouse proteomics data could provide valuable insights into the signaling networks associated with Infliximab non-response in CD.

Here, we developed an approach for translating inflammation-associated proteomics from CD mouse models to human CD Infliximab response called Translatable Components Regression (TransComp-R). TransComp-R translated proteomic insights between mouse models and humans while addressing discrepancies in molecular data types (human:transcriptomic, mouse:proteomic) and phenotypes (human:Infliximab response, mouse:inflammation). We projected a clinical cohort of human CD transcriptomic data into a mouse proteomics principal components model and performed principal components regression against the human Infliximab response phenotypes to identify the most translatable, humanized latent variables. Analysis of the proteins that defined separation along these latent variables identified activated collagen-binding integrin ITGA1 and MAP3K1 signaling that separated Infliximab responders and non-responders pre-treatment. Since CD is a complex disorder that impacts both host tissue and immune cell signaling, we obtained colonic biopsies from a CD patient and performed single-cell RNA sequencing (scRNA-seq) and analyzed publicly available datasets of FAC sorted immune cell populations to identify the cell types and inter-cellular signaling associated with the identified Infliximab resistance. From this we found that the TransComp-R identified signaling network described interactions between ITGA1+ T-cells and collagen secretion by colonic cell types and T-cell populations. Further characterization of the intercellular signaling network between immune and colonic cell types revealed a collagen binding integrin and collagen signaling network associated with Infliximab resistance in multiple disease-relevant cell types. The results of TransComp-R and our confirmatory experiments and analyses suggest that extracellular collagen and integrin signaling plays an important role in anti-TNF therapy resistance.

## Results

### The Molecular Characteristics of Infliximab Resistance Are Tissue Specific

We analyzed publicly available gene expression data of colon and ileum biopsies from Crohn’s Disease (CD) patients, prior to start of Infliximab therapy as well as 4-6 weeks after initiation of Infliximab, in comparison with healthy controls to identify signatures of Infliximab (anti-TNF) therapy resistance in each tissue (Resistant (R) versus Sensitive (S)) (2, 14). Differential expression analysis (Wilcoxon Mann-Whitney (WMW), false discovery rate correction (FDR q < 0.20)) identified several disease and Infliximab response associated-DEGs, with many more pre-treatment differences observed in colonic CD (cCD) relative to ileal CD (iCD) (Figure 1A-B). Direct comparison of ileal CD biopsies between Infliximab responders and non-responders revealed no statistically significant differences.

**Figure 1.**
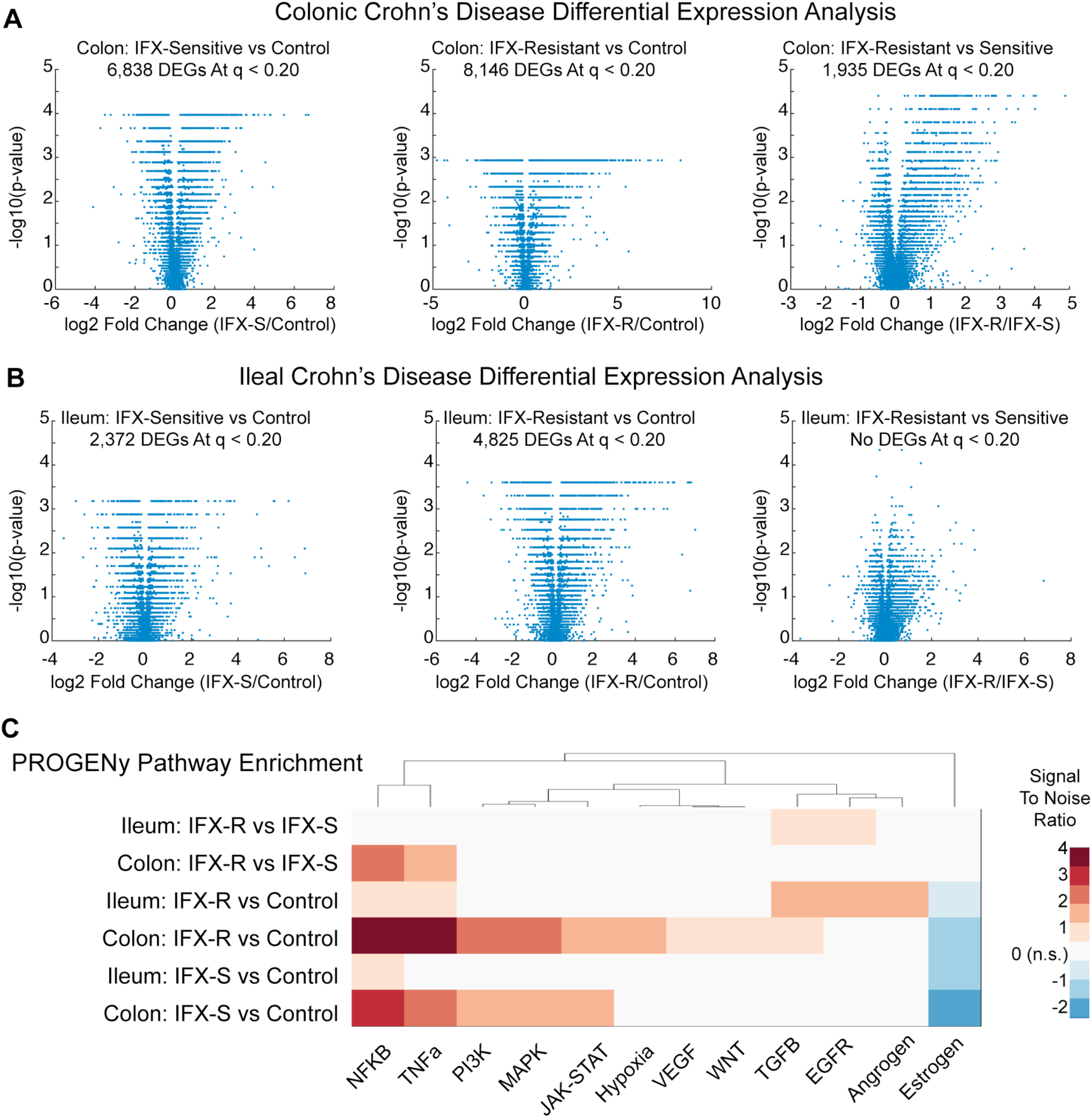
Tissue-specific signatures of Infliximab (IFX) resistance. (A) Differential expression analysis (WMW p < 0.05, FDR q < 0.20) between colonic CD Infliximab sensitive (S) (n = 12), resistant (R) (n = 7), and colon biopsies from control patients (n = 6), and (B) between ileal CD Infliximab sensitive (S) (n = 8), resistant (R) (n = 10), and ileum biopsies from control patients (n = 6). (C) PROGENy pathway enrichment analysis identified significant (WMW p < 0.05) disease and Infliximab response associated pathways.

We next assessed pathway level dysregulation between CD biopsies and controls and between Infliximab resistant and sensitive patients using PROGENy (Pathway RespOnsive Genes) (Figure 1C) (15). PROGENy infers differences in pathway activity based on high confidence signatures of downstream differentially regulated genes indicative of pathway activity rather than other approaches that use expression of pathway members to infer activity (16). Significant baseline pathway differences did exist between Infliximab sensitive and resistant patients in both cCD and iCD (WMW p < 0.05) (Figure 1C). In ileal CD, TGFB and EGFR signaling are both upregulated in Infliximab resistant patients, but these pathways were unchanged between sensitive and resistant patients with cCD. In cCD, both NFKB and TNFA are upregulated in resistant patients relative to sensitive patient, but these pathways were unchanged between sensitive and resistant patients with ileal CD. Our analysis of the colonic and ileal CD biopsies further shows that response to Infliximab is associated with tissue-specific, pre-treatment disease characteristics.

### Translatable Components Regression Predicts Proteomic Dysregulation in Patients from Mice

Though gene expression data can yield associations relevant to Infliximab responsiveness vs resistance outcomes, gene signatures alone provide an incomplete characterization of Infliximab resistance signaling in IBD (3, 4, 17). The available proteomic studies in CD have principally been serum-based and do not provide direct measurements of Infliximab resistance signaling from the site of disease (18–20). We have previously conducted high throughput proteomic measurements from two mouse models of IBD, the Adoptive T-cell Transfer (TCT) and Tumor Necrosis Factor Delta-ARE (TNF-ARE) models in inflamed and un-inflamed conditions (8, 9). However, no therapeutic stimuli were studied in these mice, which potentially limits the applicability of these datasets for understanding Infliximab resistance in patients. The evolutionary and molecular differences between mouse models and humans coupled with discrepancies in measurement types between mass-spectrometry based proteomics and microarrays further complicates translation of mouse proteomic insights to patient CD.

We developed a translational systems modeling framework called translatable components regression (TransComp-R) to address the challenges of translating mouse proteomic insights to CD patient phenotypes (Figure 2). We first trained a Principal Components Analysis (PCA) model on mouse proteomics data using proteins whose coding genes were homologs with Infliximab-response associated human DEGs. In order to link the mouse proteomic latent variables with human Infliximab response phenotypes, we needed to project our CD patient samples into mouse proteomics space. However, projecting a human transcriptomics dataset into a mouse proteomics PCA model has many analytical complications. Classically, new samples are projected into a PCA model by normalizing the new data according to the training data scaling factors and multiplying by the principal components (PC) of the PCA model. This is not appropriate for inter-species, inter-omic projections because the scaling factors cannot be assumed to be comparable between different species, sequencing platforms, or proteomic and transcriptomic data types. TransComp-R modifies the PCA projection procedure by projecting new datasets into an existing PCA model in terms of *relative differences* along mouse PCs rather than absolute differences as in the classical PCA projection procedure. To accomplish this, we projected patient samples into the mouse proteomics PCA model by multiplying the human RNA matrix by the mouse PCA eigenvectors. Once projected, we built a regression model relating human scores on mouse PCs to Infliximab response and identified the dimensions of biological variation in the mouse that best predicted Infliximab response. In a sense, the projection of human samples into mouse PC space is a form of *computational reverse-translation* and we can regard the mouse proteomic PCs that predict the human clinical associations as the most *humanized* components, or the “Translatable Components” (TCs) of the mouse (Figure 2).

**Figure 2.**
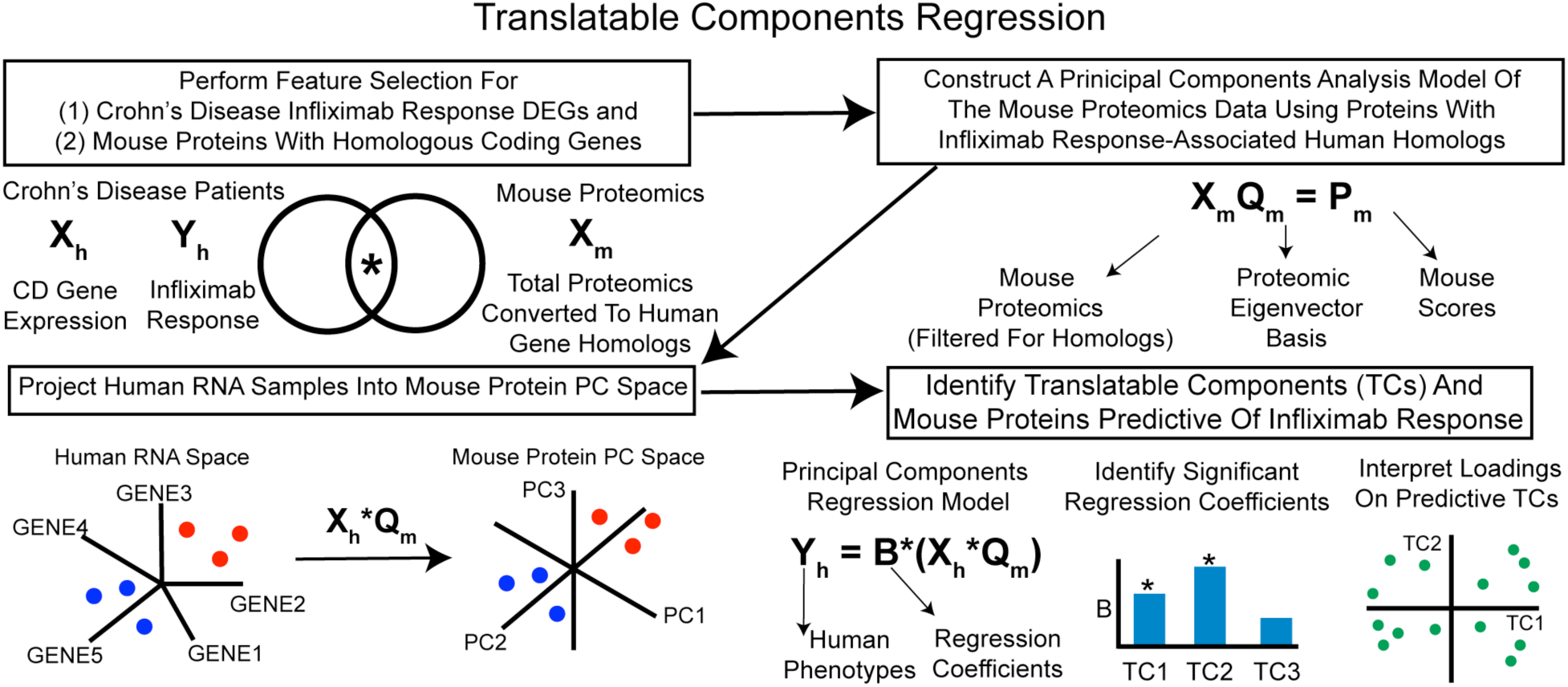
Translatable Components Regression (TransComp-R) Methodology. TransComp-R begins with feature selection of human DEGs associated with Infliximab response and mouse proteins with one-to-one human homolog DEGs. A Principal Components Analysis (PCA) model of the selected mouse proteins is then constructed. Human RNA data is projected into the mouse protein PCA model such that relative differences between human samples are represented in terms of the mouse proteomic latent variables. A Principal Components Regression (PCR) model is then constructed to relate the human scores on mouse PCs to human Infliximab response. The PCs predictive of Infliximab response (the Translatable Components (TC)) are interpreted to identify protein dysregulation associated with Infliximab response.

### TransComp-R Identifies Activated Integrin Signaling in Infliximab Resistant Colonic CD

We applied TransComp-R to identify proteomic Infliximab response signatures by incorporating CD patient pre-treatment transcriptomics data into our TCT and TNF-ARE mouse proteomics models. TCT mouse and human cCD PCA models were built using 288 of the 1,935 genes differentially expressed between cCD Infliximab responders and non-responders that had homologous protein coding mouse genes (Figure 3A). Though the Infliximab response associated genes separated the mouse samples by inflammation status, the cCD patients did not separate by Infliximab response along human PC1 and PC2. A principal components regression model (PCR) predicting Infliximab response from the cCD patient scores on mouse protein principal components (PCs) identified TCT mouse PC6 and PC7 as significantly predictive of Infliximab response (Table S1). The cCD sample scores on these PCs better separate Infliximab response than the scores on human RNA PCs, despite PC6 and PC7 accounting for only 3.3% of the variation in the TCT mouse proteomics data (Figure 3B). This contrasts with the poor cCD patient separation along mouse PC1 and PC2, which together account for 80% of the total variation in the mouse proteomics data (Figure S1). This underscores a key strength of the TransComp-R framework, the model identifies proteomic variation predictive of Infliximab response (PC6 and PC7) and ignores sources of proteomic variation that predominately associate with non-specific inflammation and inter-mouse variability (PC1 and PC2).

**Figure 3.**
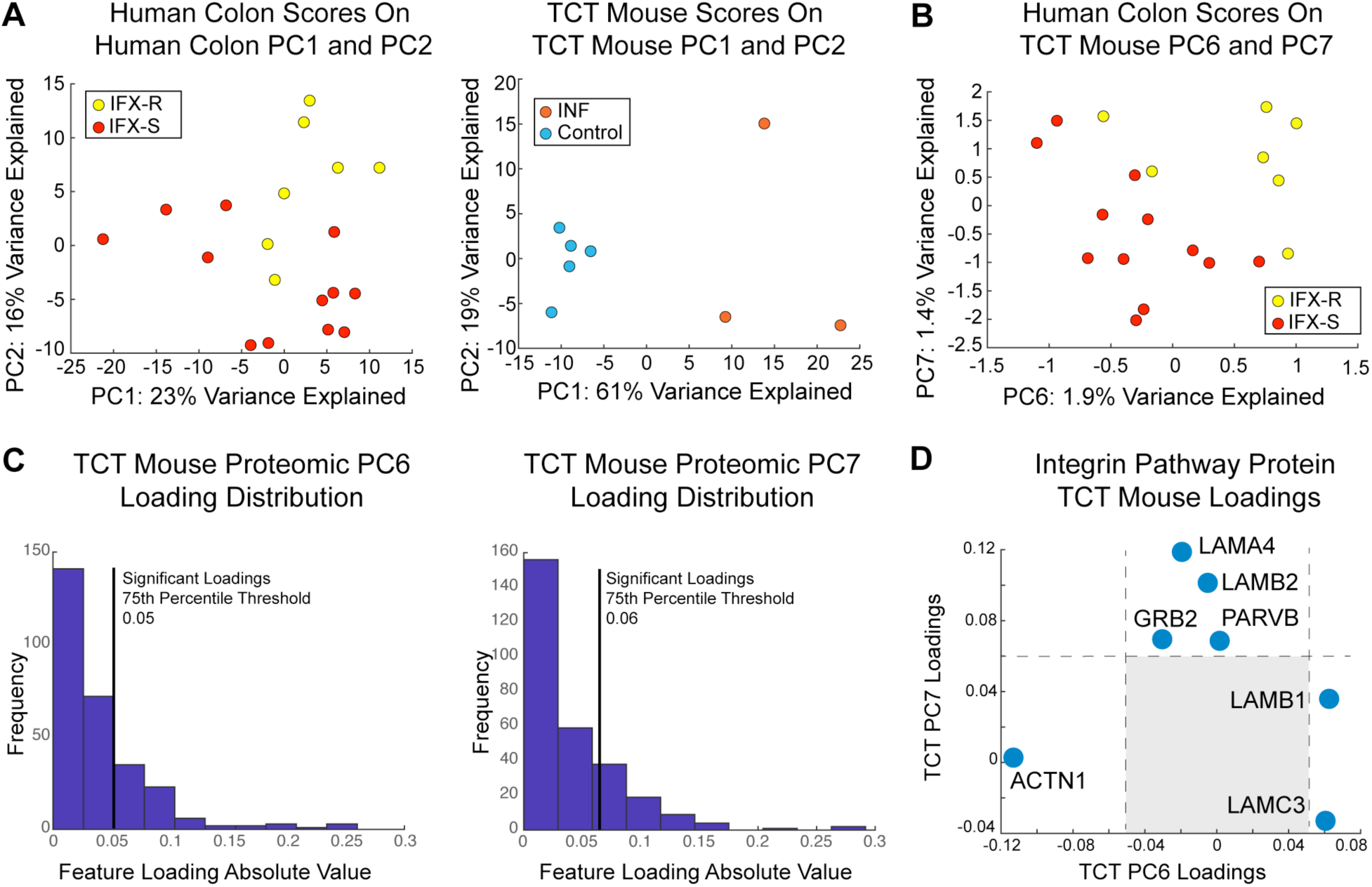
TransComp-R of cCD Infliximab response against the T-cell transfer (TCT) mouse model. (A) PCA scores plots of cCD RNA and TCT mouse proteomics data using 288 Infliximab-resistance associated differentially expressed orthologous genes. (B) TransComp-R identifies TCT mouse proteomic PC6 and PC7 as predictive of Infliximab response in cCD. (C) Histograms of absolute values of protein loadings on mouse proteomic PC6 and PC7 with 75^th^ percentile threshold for significant loadings shown for each PC. (D) Significant integrin pathway protein loadings on PC6 and PC7 with non-significant region shaded.

We plotted the empirical cumulative distribution function (eCDF) of the absolute value of the protein loadings on TCT mouse PC6 and PC7 to identify significantly loaded proteins greater than the 75th percentile of the eCDF (Figure 3C). PANTHER pathway enrichment of these proteins identified a single enriched pathway, upregulated Integrin Signaling, Infliximab resistant cCD (p = 2.68*10^-5^) (21–23). The loadings of proteins that contributed to integrin signaling enrichment were generally positive on TCT mouse PC6 and PC7, similar to the scores of colonic CD Infliximab non-responders (Figure 3D). Several laminin proteins were positively loaded toward Infliximab non-responders suggesting that interactions with the extracellular matrix (ECM) and migration-based signaling play a role in Infliximab resistance.

Since different mouse models may describe diverse aspects of disease, we performed a second TransComp-R on the cCD patients and TNF-ARE mouse proteomics data using 708 Infliximab response associated DEGs with homologous protein coding mouse genes (Figure 4A). As with the TCT mouse, the TNF-ARE mouse separated by inflammation status and the human samples did not separate well by Infliximab response on human PC1 and PC2. A PCR model predicting Infliximab response from cCD patient scores on the TNF-ARE mouse PCs identified three significantly predictive mouse PCs (Table S2). TNF-ARE mouse PC4, PC6, and PC7 were all significantly predictive of Infliximab response, with the two most predictive PC’s, PC4 and PC6, accounting for 8.9% of the total variation in the TNF-ARE proteomics. As with the TCT-mouse PCA, projecting the cCD patients into the predictive TNF-ARE PCs revealed stronger separation by Infliximab response than in the human RNA PCA or along TNF-ARE mouse PC1 and PC2 (Figure 4B, Figure S1).

**Figure 4.**
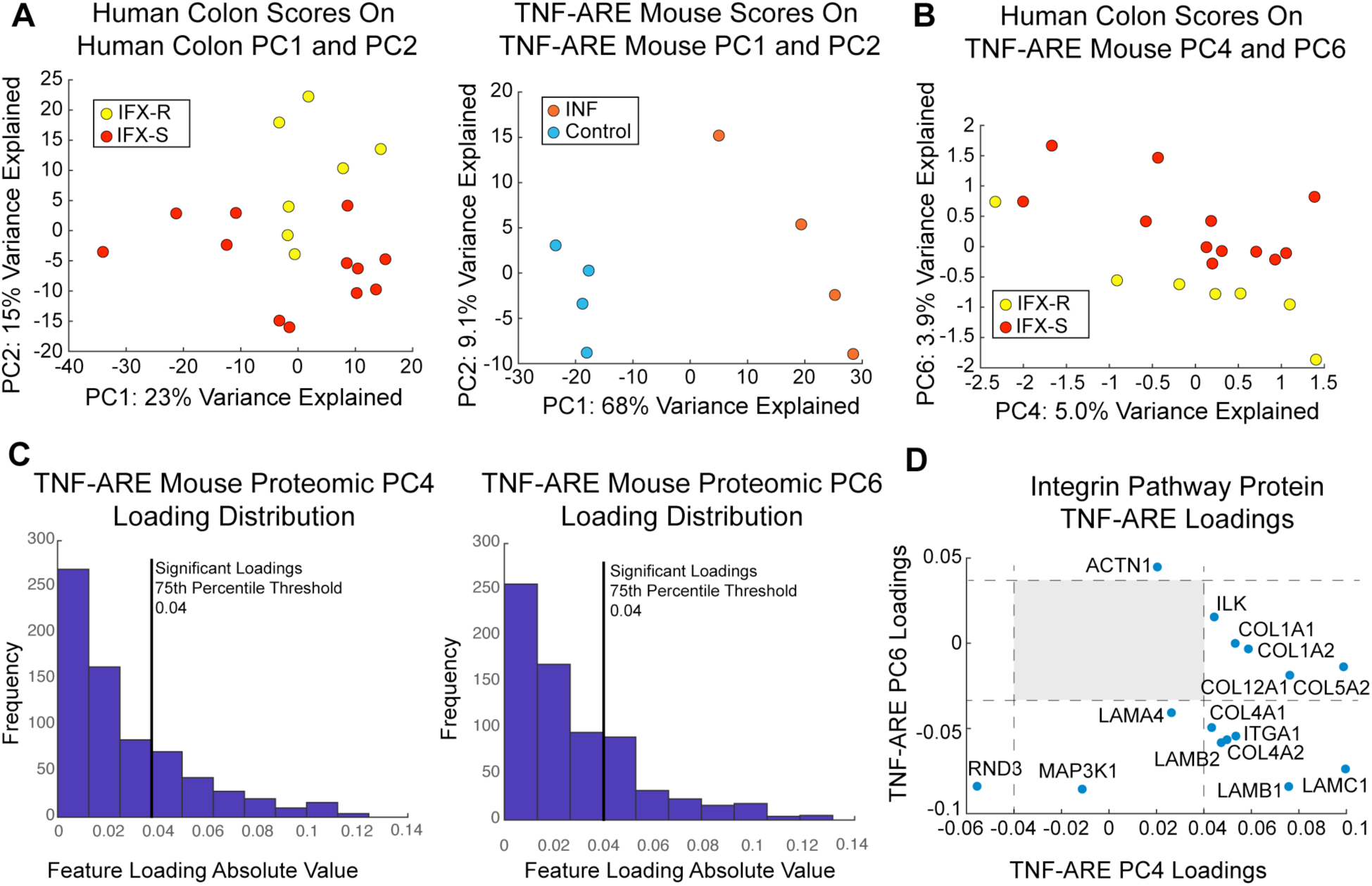
TransComp-R of colonic Crohn’s Disease Infliximab response against the TNF-ARE mouse model of IBD. (A) PCA scores plots of cCD RNA and TNF-ARE mouse proteomics data using 708 Infliximab-resistance associated DEGs with mouse homologs. (B) TransComp-R identifies TNF-ARE mouse proteomic PC4 and PC6 as significantly predictive of Infliximab response in cCD. Plotting the human scores on these mouse PCs shows stronger separation by Infliximab response than in the human RNA PCA. (C) Histograms of absolute values of protein loadings on mouse proteomic PC4 and PC6 with 75^th^ percentile threshold for significant loadings shown for each PC. (D) Significant integrin pathway protein loadings on PC4 and PC6 with non-significant region shaded.

Analysis of the protein loadings on the significant TNF-ARE PCs identified 278 significantly loaded proteins (Figure 4C). PANTHER enrichment analysis identified 20 pathways, the most significant being integrin signaling upregulation in Infliximab resistant cCD (p = 2.00*10^-5^) (Table S3). Plotting the loadings of the integrin pathway proteins once again revealed stronger weighting of integrin proteins toward Infliximab resistant cCD (Figure 4D). Laminin proteins LAMB2, LAMB1, and LAMA4 were similarly weighted toward resistant patients in both mouse models, suggesting that higher levels of laminin proteins are consistent biomarkers of Infliximab non-response translatable from either the TCT or TNF-ARE mouse models. Mitogen activated protein kinase kinase kinase 1 (MAP3K1) and integrin alpha 1 (ITGA1) were also loaded toward Infliximab resistant patients, suggesting they play a role in resistance (Figure 4D).

We found no DEGs associated with Infliximab resistance in iCD, leaving no significant features to construct a TransComp-R model. In order to see if a lower DEG significance threshold for TransComp-R could add translational value to the iCD data, we performed TransComp-R using candidate genes that had a fold change of 2 or more between Infliximab resistant and sensitive patients. We did not find any predictive TCs when applying TransComp-R to the TNF-ARE mouse proteomics data despite higher proteome coverage of human homologous genes. TransComp-R on the TCT mouse proteomics data did identify 2 significant TCs using fold change threshold of 2, but the proteins significantly loaded on these TC’s did not implicate any pathway level enrichment (Figure S2). It appears that a condition for the success of TransComp-R is the presence of a human signature of significant DEGs, but that differences in coverage of homologs were not a significant factor in performance, with the lower coverage TCT mouse data finding significant TCs with the candidate gene approach.

### Proteomic Profiling of cCD Infliximab Resistance Signaling Proteins in Immune Populations

A combination of immune and host tissue cell signaling drives Crohn’s disease pathology and determines therapeutic responsiveness. TransComp-R was performed on samples containing a mixture of colonic and immune cells making it challenging to associate Infliximab resistance signaling with particular cell types. In order to develop therapeutic strategies to overcome therapeutic resistance, it is necessary to verify the activity of the integrin signaling network in cCD patients and to characterize the cell types responsible for each component of the Infliximab resistance network. To address these questions, we performed single-cell RNA sequencing (scRNA-seq) on two biopsies from a cCD patient and analyzed a publicly available proteomics dataset of 28 FAC sorted immune cell types (ImmProt) (24).

We mined ImmProt for cell types expressing ITGA1 signaling network proteins contributing to Infliximab resistance (Figure 5) (24). Clustering of protein copy numbers revealed cell-specific high expression of certain key proteins, including specialization of MAP3K1 to neutrophils, COL4A2 to T-cell populations, ITGA1 to activated NK cells, and LAMA4 to macrophage populations (Figure 5). Other collagen and laminin proteins were moderately expressed in various populations. While broad involvement of macrophages is a hallmark of Crohn’s disease, the specificity of MAP3K1 to neutrophils suggests that this cell type is a key player in Infliximab resistance. Further, the capability of immune cell populations to secrete of collagen and laminin ligands of ITGA1 and its associated duplex integrin ITGB1 suggest a wide range of possible interactions between immune and colonic cell types may facilitate Infliximab resistance in cCD.

**Figure 5.**
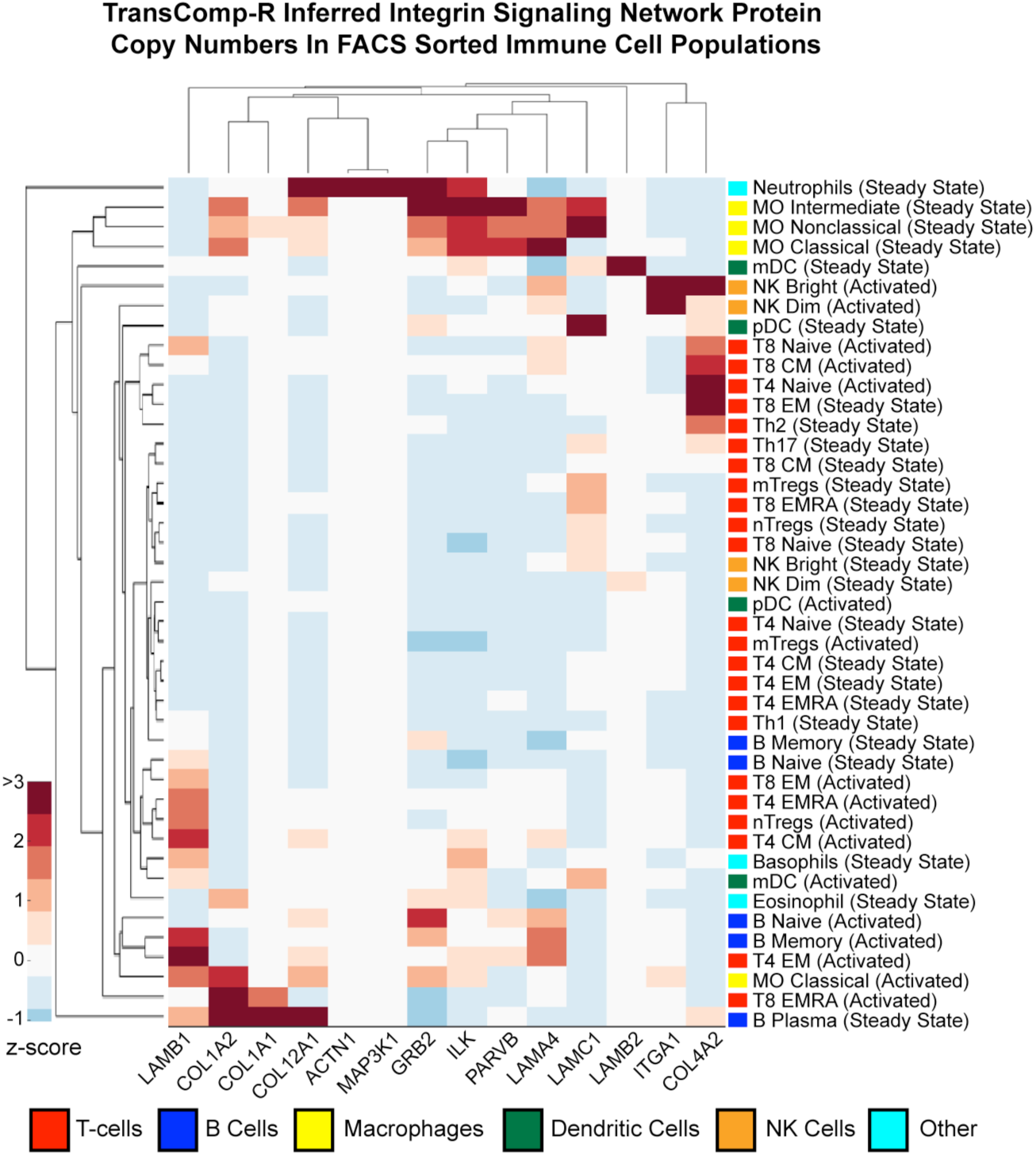
Integrin-Infliximab Resistance Network Signaling in Immune Cell Populations. Protein copy numbers from FAC sorted immune cell populations reveals cell-type specific expression of components of the Infliximab resistance network.

### Single-Cell Profiling of Infliximab Resistance-Integrin Signaling in cCD

Having shown that the Infliximab resistance associated integrin signaling network proteins were expressed in immune cell types, we next assessed ITGA1-associated signaling in the cCD context of mixed immune and colonic cell types. We first analyzed the post-Infliximab treated cCD samples from the bulk gene expression dataset to see if dysregulation in ITGA1-associated signaling persisted after treatment. Both before and after treatment, the Infliximab resistance-associated integrin pathway proteins were more highly expressed in resistant patients relative to sensitive patients (Figure S3). Activity of this signaling network after treatment suggests that the ITGA1 signature is a durable feature of Infliximab resistant cCD biology that is suppressed, but not resolved, by Infliximab treatment. We could therefore compare the activity of the ITGA1 signaling network from TransComp-R analysis to scRNA-seq data from left and right colonic biopsies obtained from a CD patient post-anti-TNF treatment and characterize the cell types and inter-cellular signaling network of ITGA1-associated Infliximab resistance.

Cells were filtered for those expressing at least one of the 18 genes in the Infliximab resistance signature. The right colon biopsy was filtered from 3,922 cells to 2,153 cells and the left colon filtered from 1,329 cells to 903 cells. We applied a Gaussian Mixture Model (GMM) based approach to classify cell types in each biopsy using a set of marker genes curated from the literature (Table S4) (25–27). The GMM identified four distinct cell types including a general epithelial cell population, goblet cells, stromal cells, and T-cells in each biopsy which we visualized using t-distributed stochastic neighbor embedding (TSNE) (Table S5, Figure 6A-B). Though it is possible other sub-populations of cell types were present within the large epithelial category, the accuracy of the GMM classification degraded on our datasets when we tried to predict additional cell types.

**Figure 6.**
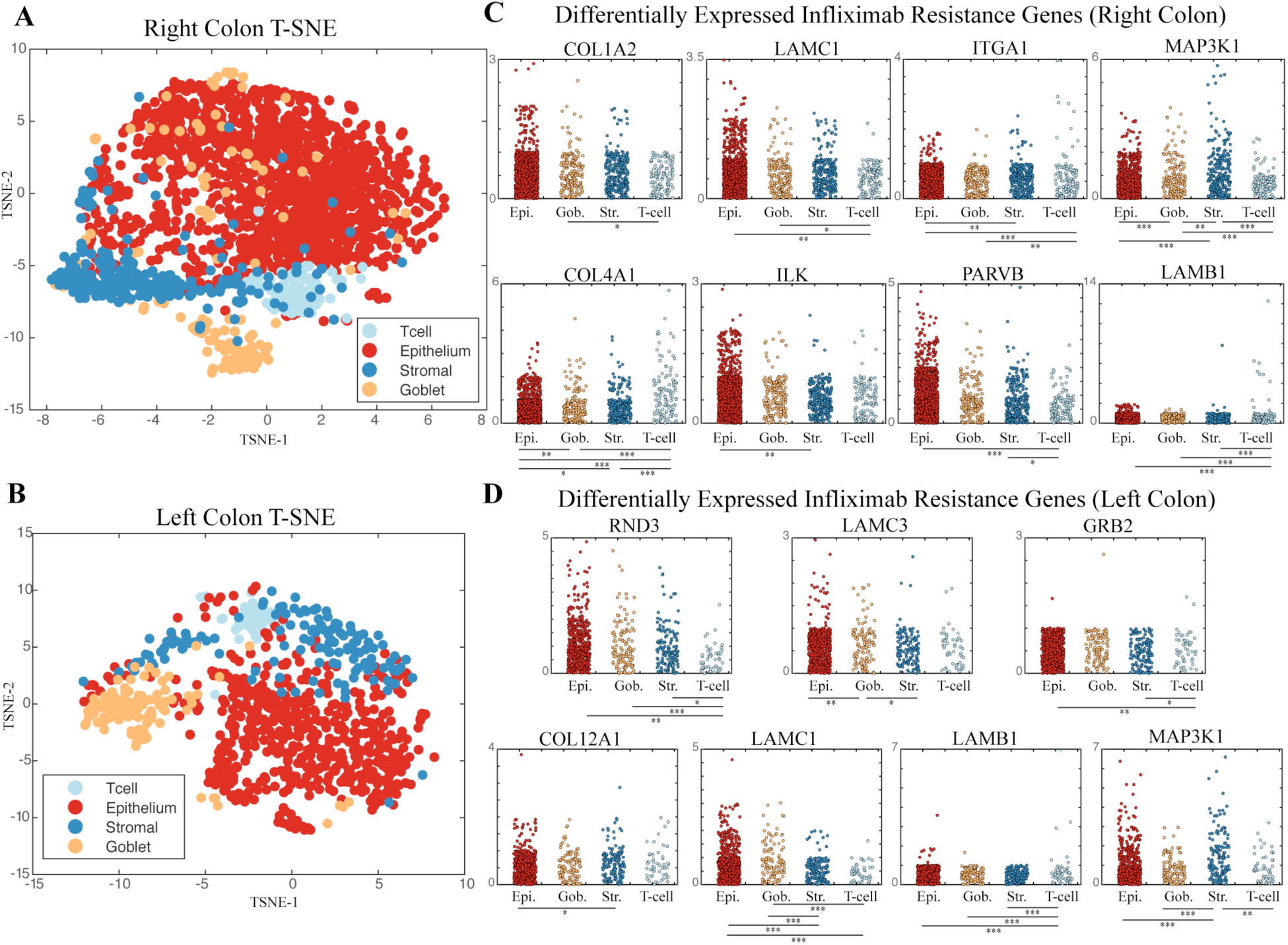
Single-cell analysis of Infliximab resistance network genes in colonic CD. (A) TSNE plot of cells expressing resistance markers from right colonic CD biopsy. (B) TSNE plot of cells expressing resistance markers from left colonic CD biopsy. (C) Differentially expressed (Kruskal-Wallis p< 0.05, WMW p < 0.05) resistance markers in right colonic CD biopsy. (D) Differentially expressed resistance markers in left colonic CD biopsy.

We performed differential expression analysis (Kruskal-Wallis test) on the Infliximab resistance signature genes to characterize cell type specific signaling of the ITGA1-associated signaling network (Table S6, Figure 6C-D). Eight genes were differentially expressed between cell types in right colonic CD (ITGA1, MAP3K1, LAMB1, LAMC1, ILK, COL1A2, COL4A1, PARVB) and seven genes were differentially expressed in left colonic CD (MAP3K1, LAMB1, LAMC1, RND3, GRB2, COL12A1, LAMC3). Three genes (MAP3K1, LAMB1, LAMC1) were similarly differentially expressed between cell types in both biopsies, with MAP3K1 more highly expressed in stromal cells, LAMB1 overexpressed in T-cells, and LAMC1 underexpressed in T-cells (Figure 6C-D). ITGA1 was significantly overexpressed in T-cells relative to colonic cell types in right colonic CD and highly expressed in T-cells and goblet cells in left colonic CD, though the differences in left colonic CD were not significant.

### Characterizing Infliximab Resistance Signaling Between ITGA1+ T-cells Colonic Cell Types

Having identified a population of ITGA1+ T-cells as potential mediators of Infliximab resistance, we next characterized communication between T-cells and colonic cell types using a single-cell inter-cellular scoring algorithm that identifies active, potentially targetable, ligand-receptor (LR) interactions between cells (25). We scored 2,567 receptor-ligand interactions and identified 16 significant interactions (top 10% of all interaction scores) in left colonic CD containing at least one Infliximab resistance signature gene, all of which were also present in the 22 significant interactions in right colonic CD.

We visualized these interactions between cell types in a network using Cytoscape, with nodes for each cell type and edges indicating if a significant LR interaction was present colored by the cell expressing the receptor or ligand (Figure 7A) (28). T-cells predominately interacted with each other through collagen COL1A1 binding to integrin ITGB1, the other half of the integrin α1β1 duplex with ITGA1 (Figure 7B). ImmProt analysis showed that ITGB1 protein was highly abundant in all ImmProt T-cell populations, but the relative specificity of ITGA1 and COL1A1 in different T-cell populations suggests that these proteins interactions with ITGB1 determine the specificity of T-cell to T-cell interactions in cCD (Figure 7B) (24).

**Figure 7.**
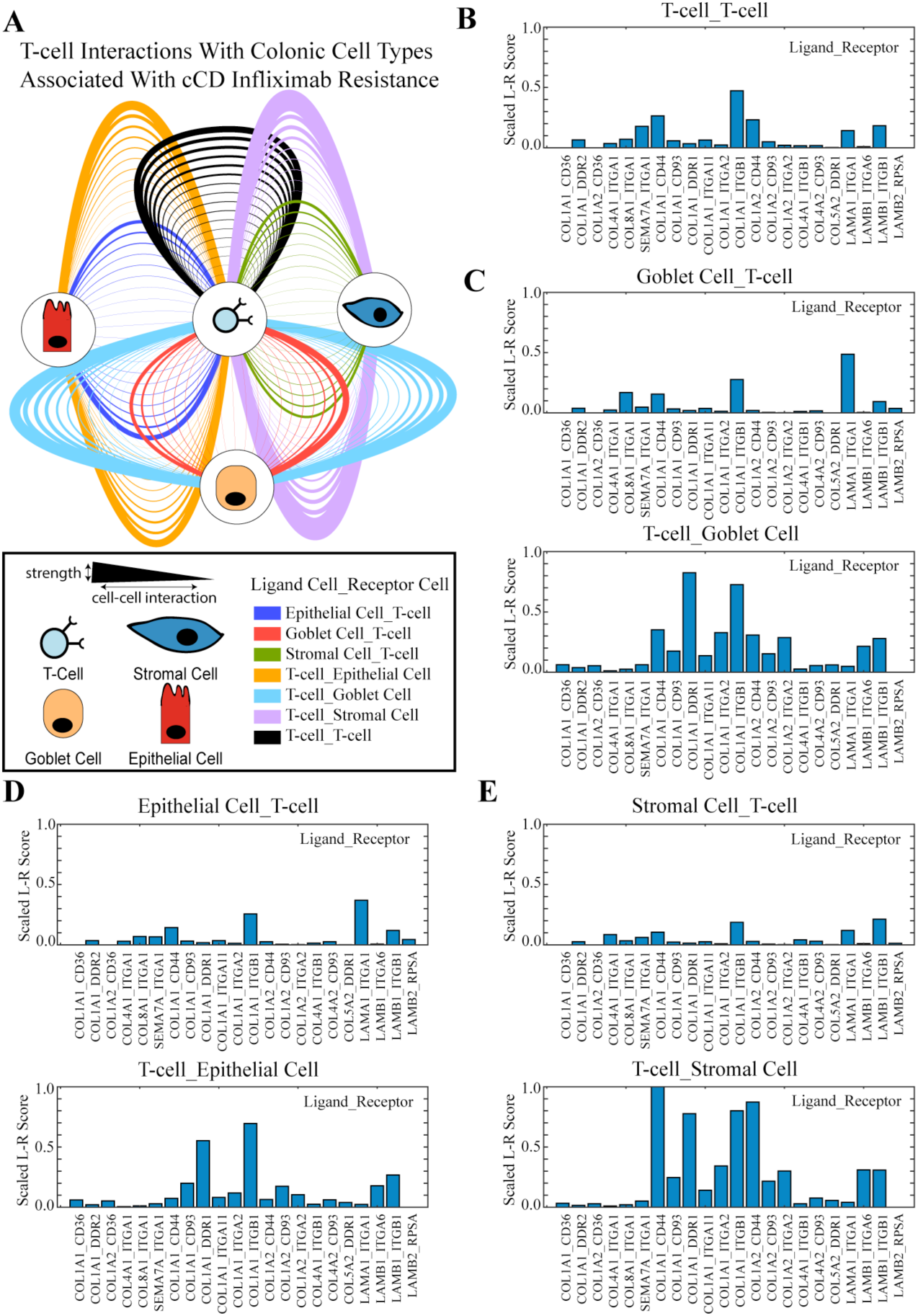
Cell-cell ligand-receptor interaction scoring of Infliximab resistance associated genes in cCD. (A) Network of significant ligand-receptor (LR) scores containing at least one Infliximab resistance signature gene. Cell types are connected by edges indicating a LR interaction was significant present and edge thickness indicates the strength of interaction. (B) Normalized LR scores for T-cell to T-cell interactions. (C) Normalized LR scores for T-cell and goblet cell interactions. (D) Normalized LR scores T-cell and epithelial cell interactions. (E) Normalized L-R scores for T-cell and stromal cell interactions.

Interactions via the integrin α1β1 duplex were highly prevalent between ITGA1 positive T-cells and colonic cell types (Figure 7C-E). ITGA1 positive T-cells predominately interact with goblet cells via goblet cell secretion of LAMA1 interacting with ITGA1 or COL1A1 interacting with ITGB1, the other half of the integrin α1β1 duplex (Figure 7C). Goblet cells in turn appeared to express the integrin α2β1 duplex and bind to COL1A1 secreted by T-cells, but the strongest interaction was between COL1A1 secreted by T-cells and the discodin domain receptor tyrosine kinase, DDR1, on goblet cells (Figure 7C). T-cell interactions with other epithelial cell populations were mostly mediated by collagens secreted by T-cells binding to either integrin α1β1 or integrin α2β1 or DDR1 (Figure 7D). ITGA1 and ITGB1 expressing T-cells primarily interacted with epithelial cells by binding to secreted COL1A1 or laminins LAMA1 or LAMB1 (Figure 7D).

Interactions between T-cells and stromal cells accounted for the highest number of significant interactions (Figure 7E). The interaction between COL4A1 secreted by stromal cells and ITGA1+ T-cells was stronger here than between other pairs of cell types and was the only significant interaction that contained two proteins from our Infliximab resistance signature. We also observed secretion of COL1A1 by T-cells and subsequent binding to DDR1, ITGB1, and ITGA2 accounted for the most significant cell-cell interactions across all cell types (Figure 7). The large number of significant interactions between ITGA1+ T-cells and colonic cell types and large number of interactions involving COL1A1 secretion by T-cells suggest that collagen binding integrins, expressed on both T-cells and colonic cell types, play a role in Infliximab resistance and that the interactions between these cell types may be potential therapeutic targets.

## Discussion

The broadly important, fundamental challenge of inter-species translation can often become further complicated by discrepancies in phenotypes and molecular data types between clinical cohorts and pre-clinical experimental systems. Here, we demonstrated the effectiveness of the TransComp-R framework for translating proteomic dysregulation in CD mouse models a phenotypically and molecularly mismatched clinical cohort of CD patients to characterize resistance to anti-TNF therapy. We identified a network signature for Infliximab resistance that links many disparate observations of laminins, collagen binding integrins, and MAP3K1 signaling in cCD to the clinically important challenge of overcoming resistance to anti-TNF therapies. Our verification of the relevance of this signature in immune cell proteomics data and in independent biopsies from a cCD patient indicates a larger role for collagen, laminin, and associated integrin binding signaling in CD pathobiology and to a particular therapeutic resistance phenotype.

Overall, our results indicate an expanded role for collagen binding integrin signaling in the clinically important phenotype of Infliximab resistant colonic Crohn’s Disease. In healthy colon, ITGA1 expression localizes to colon crypts in (29) and inhibition of ITGA1 is protective against colitis in the DSS and TNBS mouse models of IBD (30, 31). Previous studies examining tissue-independent markers of memory T-cells have also implicated ITGA1 as a consistent surface marker of tissue-resident memory T-cells (32). Our results suggest that collagen binding integrin signaling associated with Infliximab resistance is facilitated predominately by a memory T-cell population and their interactions with other colonic cell types. The combined evidence of our ligand-receptor scoring of our scRNA-seq data and the ImmProt analysis indicates that the T-cells expressing ITGA1 may not be the same one secreting collagen and laminin proteins into the extracellular space, that other populations of T-cells secrete ligands eliciting integrin signaling in memory T-cell and colonic cell populations. The downstream consequences of this complex intercellular signaling network may in turn activate intracellular pathways on T-cells and colonic cell types that facilitate CD disease progression and result in Infliximab resistance.

Our results also indicate that MAP3K1 signaling may be playing a role in Infliximab resistance as an intracellular mediator of the extracellular signaling cues from collagen binding integrin signaling. Signaling to ITGA1 on the surface of cells is capable of regulating MAP3K1 via GIRB, another Infliximab resistance protein identified by TransComp-R (33–36). Further studies have shown that MAP3K1 is a regulator of RAF1, itself a mediator of the ERK and JNK signaling cascades (37). A small clinical trial of a RAF1 inhibitor showed that targeting the JNK signaling cascade in macrophages could induce remission in cCD and that this could be achieved in Infliximab non-responsive patients (38–40). MAP3K1 also carries a colonic CD specific SNP, rs832582, may predispose carriers to a more intense inflammatory response (41). A potential biomarker of Infliximab resistance in cCD could therefore be MAP3K1 signaling in neutrophils and/or stromal cells as well as potentially the IBD risk SNP rs832582 for MAP3K1. Importantly, the convergence of multiple signaling disruptions by collagen binding integrins and MAP3K1 suggests that patients expressing these disease characteristics would not benefit from anti-TNF therapy and should be considered for an alternative therapeutic course.

A powerful computational feature of TransComp-R is that it identifies mouse proteomic PCs predictive of human phenotypes despite these PCs explaining little variation in the mouse proteomics data. In standard PCA, latent variables are constructed to explain the variance in the training dataset (mouse proteomics), rather than the relationship of the PCs to a phenotype or to reflect variance of a secondary dataset (human transcriptomics). Since the mouse PCA model is built using mice with different phenotypes than the human CD dataset (mouse inflamed vs. uninflamed, human Infliximab responder vs. non-responder), it is not surprising that mouse PC1 and PC2 do not separate projected human phenotypes. This suggests that the most translatable pre-clinical biology may not be that which most immediately explicates the experimental groups, but instead indicates that a computational modeling approach such as TransComp-R can more insightfully recover translationally relevant biology (for instance, as obscured in less obvious PCs). We believe that Translatable Components Regression is widely applicable to challenges of translation in other disease contexts, model systems, and types of molecular data.

## Materials and Methods

### Analysis of Human CD Gene Expression Data

Colonic and ileal CD transcriptomic data was obtained from gene expression omnibus, accession number GSE16879, using Bioconductor tools and normalized by robust multichip average method (2, 14, 42, 43). Differential expression analysis was performed using the Wilcoxon Rank Sum test with Benjamini Hochberg False Discovery Rate (FDR) correction and significance defined by q < 0.20. PROGENy pathway enrichment analysis was run on the human data as previously described (15). Pathways were tested for differential regulation by Wilcoxon Rank Sum test with p < 0.05 considered significant.

### Analysis of Mouse Proteomics Datasets

The T-cell transfer and TNF-ARE mouse proteomics datasets were obtained from two studies examining proteomic changes between inflamed and uninflamed mice (8, 9). The mouse protein identifiers were mapped to their coding genes and converted to human gene symbols using the Mouse Genome Informatics databases (44, 45). Only one-to-one mouse-human homologs were retained for the analysis.

### Translatable Components Regression

When constructing a PCA model, it is often desirable to project observations from another dataset into that model to examine how the variability explained by the model relates to those new observations. This requires normalizing the new observation, usually by mean centering and scaling by the standard deviation of the data used to train the PCA model. However, if the new observation is measured on a different sequencing platform, comes from a different species, or is of a different molecular data type, then this centering and scaling by training data factors is not well defined and may distort the projected observation. Therefore, cross-species, cross-omic, and cross-platform projections of biological datasets and observations should not be undertaken by the standard PCA projection method. The primary component of a PCA model is identification of the eigenvectors of the covariance matrix of the training data, that is, the PCs that explain the greatest possible amount of variability in the training dataset. Though these vectors define a basis that has a particular interpretation for the training dataset, we can ask how new observations project *relative* to this coordinate system. This is done by first internally normalizing the new observations by their own mean and standard deviation to define the relative spread of each variable and then multiplying these normalized observations by the eigenvectors of the training dataset. Once projected, we performed principal components regression of the projected data against any outcome or phenotypic variable of the new observations to identify the PCs of the training data that best explain the phenotype from the new observations.

### Immune Cell Proteomics Analysis

FAC sorted quantitative proteomics data was obtained from (24) and analyzed for protein copy numbers of significantly loaded integrin pathway proteins identified by TransComp-R (Figure 5). Immune populations not expressing any protein from the network were excluded along with proteins not measured in the dataset. Data were z-score normalized by protein and clustered to identify groups of co-regulated proteins expressed in similar immune cell types. Analysis was performed in MATLAB_R2018b.

### Collection of Patient Samples

The study protocol was approved by the Institutional Review Board at Vanderbilt University Medical Center. Written informed consent was obtained for analyses of analysis of demographics, medication history, serum, and tissue biopsies obtained at the time of endoscopic procedures as part of routine clinical care to evaluate for disease activity and response to therapy in a patient with ileo-colonic CD. The patient underwent an overnight fast and received polyethylene glycol electrolyte solution for bowel preparation prior to colonoscopy. At the time of colonoscopy, biopsy specimens were obtained from the right and left colon (2 bites in each location with large capacity biopsy forceps). The specimens were placed in a 1.5mL Eppendorf tube with RPMI media, placed on ice, and transported to the lab for further processing for scRNA-seq analysis.

### Tissue processing

Biopsies were delivered from endoscopy in cold RPMI, and transferred to DPBS (without Ca or Mg) with 4mM EDTA and .5mM DTT to chelate for one hour before being lightly triturated in DPBS. Tissues were then resuspended DPBS containing cold-active protease (Sigma) at 5mg/ml with DNase (Sigma) at 2.5mg/ml and incubated for 20 minutes at 4-6°C with rocking motion. Trituration with a P1000 pipette needle was performed on dissociated suspension to yielded single cells, which were then filtered through a 35µm mesh and washed into DPBS. Essentially, the entire specimen was dissociated and used for subsequent steps. Live cell concentration was counted based on Trypan Blue positive cells. Cells were adjusted to a concentration of 150,000 cells/ml and Optiprep was added to a final concentration of 16% just prior to encapsulation.

### inDrop single-cell RNA-seq

Single-cell encapsulation of gut epithelial tissue was performed using the inDrop platform (1CellBio) with an in vitro transcription library preparation protocol, as previously described (46, 47). inDrop utilizes CEL-Seq in preparation for sequencing and is summarized as follows: (1) reverse transcription (RT), (2) ExoI nuclease digestion, (3) SPRI purification (SPRIP), (4) second strand synthesis, (5) SPRIP, (6) T7 in vitro transcription linear amplification, (7) SPRIP, (8) RNA fragmentation, (9) SPRIP, (10) primer ligation, (11) RT, and (12) library enrichment PCR. Number of cells encapsulated was calculated by approximating the density of single-cell suspension multiplied by bead loading efficiency during the duration of encapsulation. Approximately 3,000 cells for each sample entered the microfluidic chip. Following library preparation, as described above, the samples were sequenced using Nextseq 500 (Illumina) using a 150 bp paired-end sequencing kit in a customized sequencing run. After sequencing, reads were filtered, sorted by their designated barcode, and aligned to the reference transcriptome using the InDrop pipeline (48, 49).

### Single-cell filtering

scRNA-seq count data was filtered using several steps. First, the cumulative read inflection point was plotted. A cutoff of approximately 25-30% beyond the inflection point was used to exclude low quality barcodes, but retain cells with small library sizes. The filtered data were then normalized for library size and transformed, and visualized using t-SNE with 100 PCs. Density peak clustering was performed, and user-defined thresholds were set to obtain 10-20 clusters. Library size rank and combined mitochondrial gene expression were overlaid onto t-SNE space and low-quality cells were removed using these criteria. Canonical marker genes of cell types were also overlaid onto the t-SNE space to ensure that cell types of interest were not removed during filtering.

### Cell Type Classification and Ligand-Receptor Interaction Scoring

Cells were filtered for subsets expressing at least Infliximab resistance marker gene from the TransComp-R signature. Cell type markers were selected from previous single-cell analyses of colonic and intestinal tissue contexts and used to train a Gaussian mixture model (GMM) on the log-normalized expression data as previously described (25–27). Differential expression analysis was carried out using the Kruskal-Wallis test (p< 0.05) on Infliximab resistance marker genes. We then characterized the intercellular signaling network of ligand-receptor interactions between identified cell types by assigning a score based on the product of average receptor expression in a cell type with the average ligand expression in the interacting cell type as previously described (25). Receptor-ligand interaction scores in the top 10% of all interaction scores across cell types that contained at least one Infliximab resistance signature gene were retained for downstream analysis and interpretation.

## Supplementary Information

**Fig. S1.**
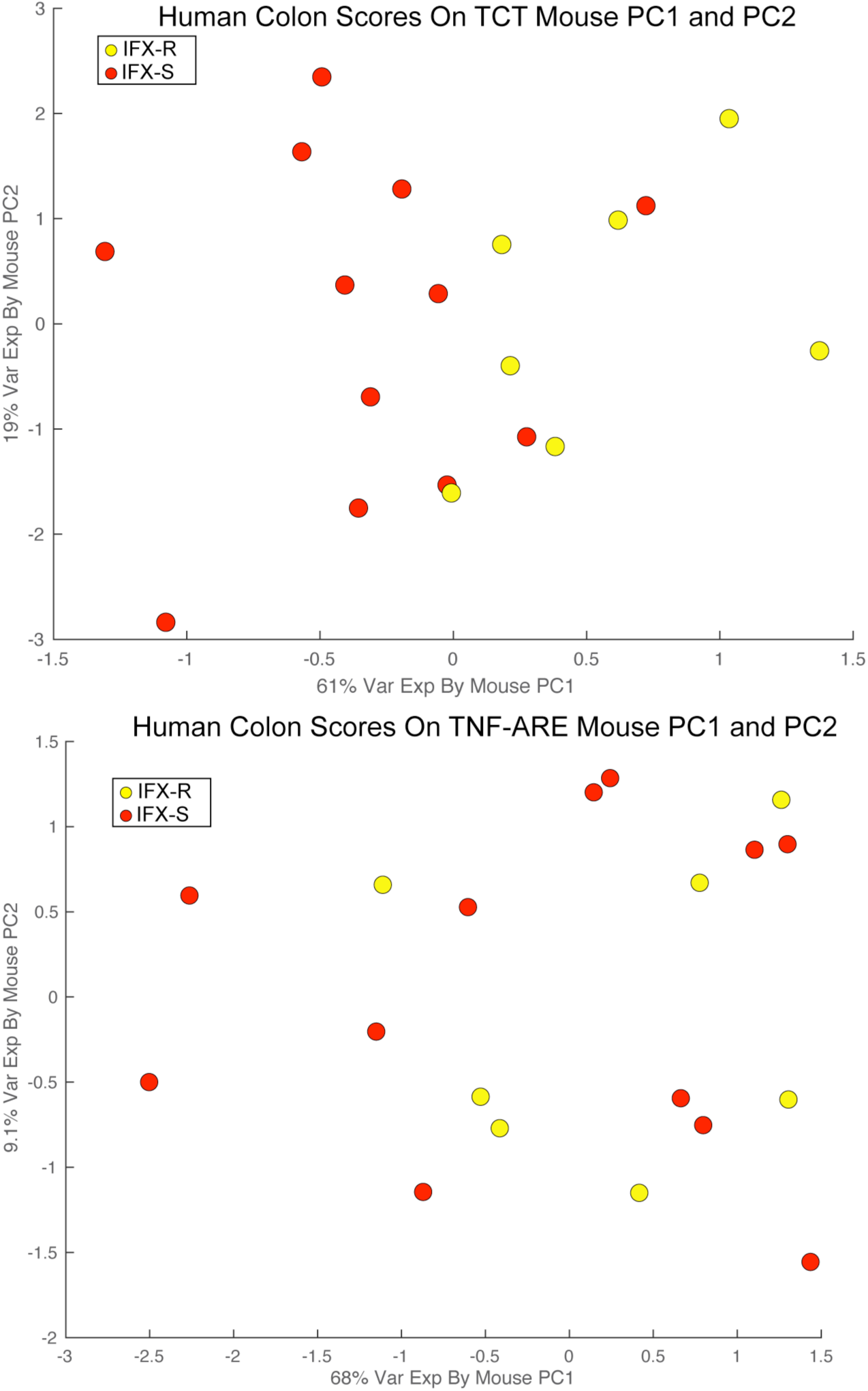
cCD patient scores on non-predictive mouse PCs. Human cCD patient scores on (A) TCT mouse proteomic PC1 and PC2 and (B) TNF-ARE mouse proteomic PC1 and PC2.

**Fig. S2.**
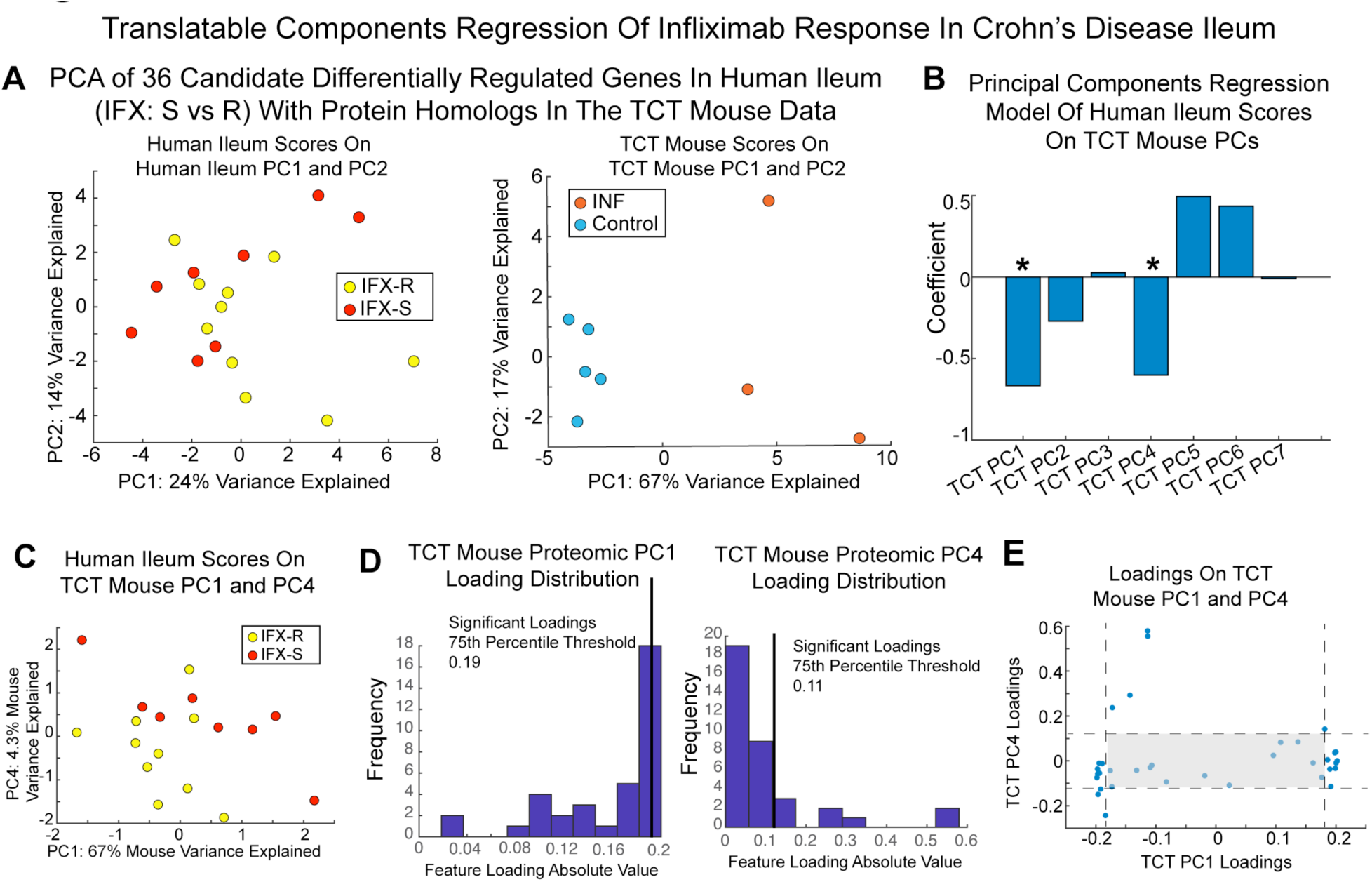
TransComp-R of iCD patient and TCT mouse data. TransComp-R of iCD and TCT mouse proteomics (A) iCD scores on human RNA PC1 and PC2 and T-cell transfer (TCT) mouse scores on mouse proteomic PC1 and PC2. (B) PCR model coefficients. (C) Human scores on mouse proteomic PC1 and PC4. (D) Distribution of absolute values of protein loadings on mouse proteomic PC1 and PC4. (E) Loadings plot with non-significant protein loading bounds shaded in grey.

**Fig. S3.**
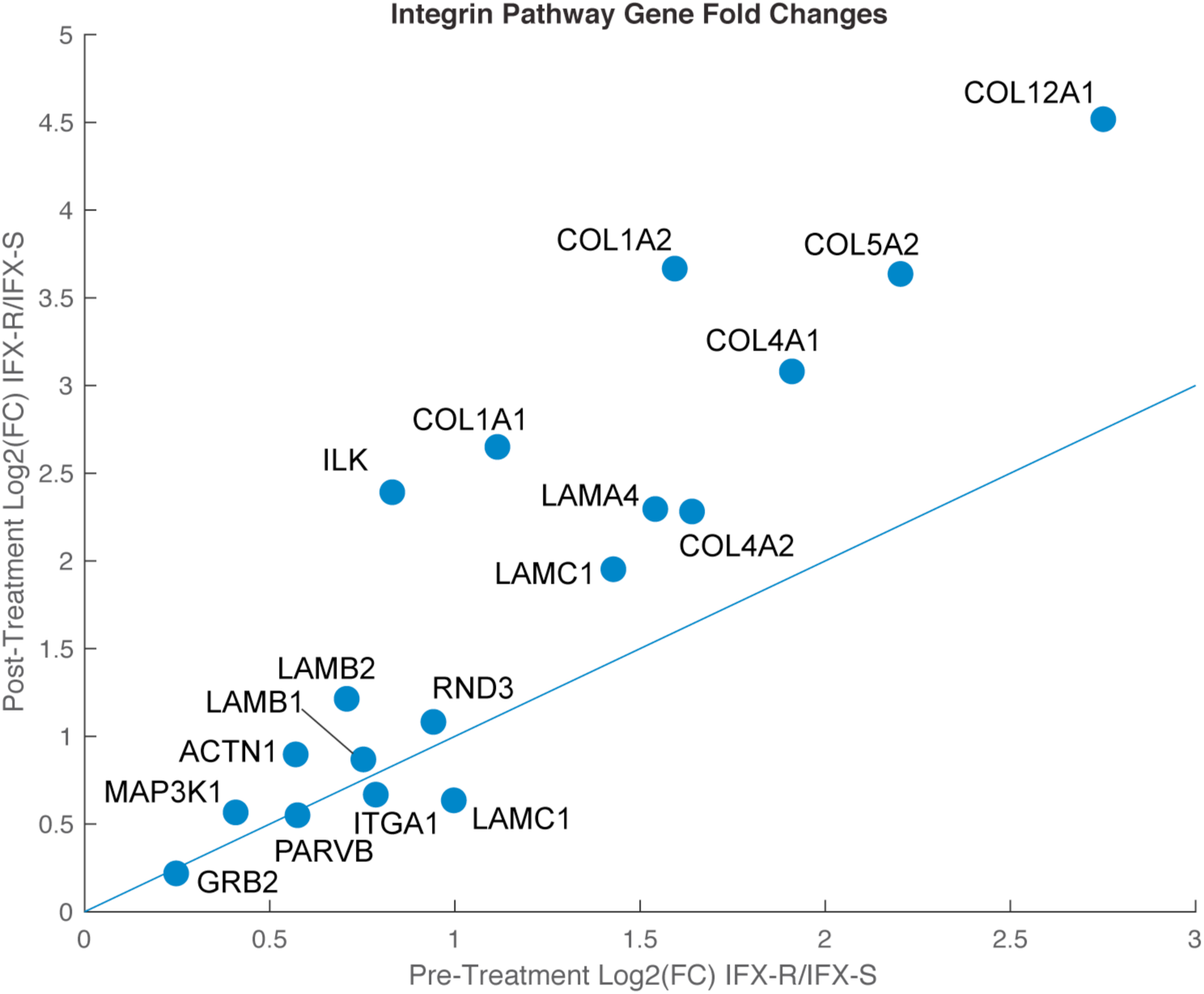
Comparison of integrin pathway gene expression between resistant (R) and sensitive (S) patients before and after Infliximab treatment.

**Table S1.** Regression coefficients for the T-cell Transfer mouse and colonic Crohn’s Disease TransComp-R model.

| Model Coefficient | Estimate | SE | tStat | pValue |
| --- | --- | --- | --- | --- |
| Intercept | -0.26 | 0.11 | -2.46 | 0.032 |
| Mouse Proteomic PC1 | 0.37 | 0.25 | 1.44 | 0.18 |
| Mouse Proteomic PC2 | -0.07 | 0.12 | -0.60 | 0.56 |
| Mouse Proteomic PC 3 | 0.023 | 0.14 | 0.15 | 0.88 |
| Mouse Proteomic PC 4 | 0.27 | 0.21 | 1.29 | 0.22 |
| Mouse Proteomic PC 5 | 0.10 | 0.13 | 0.81 | 0.43 |
| Mouse Proteomic PC 6 | 1.04 | 0.26 | 3.91 | 0.0024 |
| Mouse Proteomic PC 7 | 0.33 | 0.13 | 2.63 | 0.023 |

**Table S2.** Regression coefficients for the TNF-ARE mouse and colonic Crohn’s Disease TransComp-R model.

| Model Coefficient | Estimate | SE | tStat | pValue |
| --- | --- | --- | --- | --- |
| Intercept | -0.26 | 0.13 | -1.96 | 0.076 |
| Mouse Proteomic PC1 | -0.10 | 0.18 | -0.58 | 0.57 |
| Mouse Proteomic PC2 | 0.21 | 0.21 | 1.03 | 0.33 |
| Mouse Proteomic PC3 | -0.17 | 0.14 | -1.21 | 0.25 |
| Mouse Proteomic PC4 | -0.49 | 0.17 | -2.89 | 0.015 |
| Mouse Proteomic PC5 | -0.082 | 0.20 | -0.41 | 0.69 |
| Mouse Proteomic PC6 | -0.96 | 0.23 | -4.27 | 0.0013 |
| Mouse Proteomic PC7 | 0.39 | 0.14 | 2.85 | 0.016 |

**Table S3.** PANTHER pathway enrichment analysis of significantly loaded proteins on TNF-ARE mouse PC4 and PC6.

| PANTHER Pathways | p-value | FDR q-value |
| --- | --- | --- |
| Integrin signaling pathway (P00034) | 1.22E-07 | 2.00E-05 |
| Histamine H1 receptor mediated signaling pathway (P04385) | 4.72E-05 | 1.93E-03 |
| Muscarinic acetylcholine receptor 1 and 3 signaling pathway (P00042) | 2.85E-05 | 2.33E-03 |
| Corticotrophin releasing factor receptor signaling pathway (P04380) | 1.12E-04 | 3.66E-03 |
| Angiotensin II-stimulated signaling through G proteins and beta-arrestin (P05911) | 2.36E-04 | 4.83E-03 |
| Oxytocin receptor mediated signaling pathway (P04391) | 1.96E-04 | 5.36E-03 |
| Thyrotropin-releasing hormone receptor signaling pathway (P04394) | 2.33E-04 | 5.46E-03 |
| Heterotrimeric G-protein signaling pathway-Gq alpha and Go alpha mediated pathway (P00027) | 4.34E-04 | 7.11E-03 |
| 5HT2 type receptor mediated signaling pathway (P04374) | 4.08E-04 | 7.43E-03 |
| Beta3 adrenergic receptor signaling pathway (P04379) | 8.43E-04 | 1.26E-02 |
| Opioid proenkephalin pathway (P05915) | 1.47E-03 | 1.61E-02 |
| 5HT4 type receptor mediated signaling pathway (P04376) | 1.32E-03 | 1.67E-02 |
| Gonadotropin-releasing hormone receptor pathway (P06664) | 1.64E-03 | 1.68E-02 |
| Opioid proopiomelanocortin pathway (P05917) | 1.47E-03 | 1.72E-02 |
| Opioid prodynorphin pathway (P05916) | 1.32E-03 | 1.81E-02 |
| Muscarinic acetylcholine receptor 2 and 4 signaling pathway (P00043) | 2.04E-03 | 1.97E-02 |
| 5HT1 type receptor mediated signaling pathway (P04373) | 4.16E-03 | 3.41E-02 |
| Beta1 adrenergic receptor signaling pathway (P04377) | 4.16E-03 | 3.59E-02 |
| Metabotropic glutamate receptor group II pathway (P00040) | 4.80E-03 | 3.75E-02 |
| Beta2 adrenergic receptor signaling pathway (P04378) | 4.16E-03 | 3.79E-02 |

**Table S4.** Marker genes for classification of cell types in the colonic scRNA-seq datasets.

| Biopsy | Cell Type | Marker Gene |
| --- | --- | --- |
| Left | T-cell | CD2 |
| Left | T-cell | TRAC |
| Left | T-cell | TRBC2 |
| Left | T-cell | CD3D |
| Left | Epithelium | EPCAM |
| Left | Epithelium | KRT19 |
| Left | Epithelium | KRT20 |
| Left | Goblet | EPCAM |
| Left | Goblet | MUC2 |
| Left | Goblet | REG4 |
| Left | Stromal | RPS27 |
| Left | Stromal | RPS12 |
| Left | Stromal | RPS24 |
| Left | Stromal | SLC12A2 |
| Right | T-cell | CD2 |
| Right | T-cell | TRAC |
| Right | T-cell | TRBC2 |
| Right | Epithelium | EPCAM |
| Right | Epithelium | KRT19 |
| Right | Goblet | CLCA1 |
| Right | Goblet | MUC2 |
| Right | Goblet | REG4 |
| Right | Stromal | RPS27 |
| Right | Stromal | RPS12 |
| Right | Stromal | RPS24 |
| Right | Stromal | SLC12A2 |

**Table S5.**
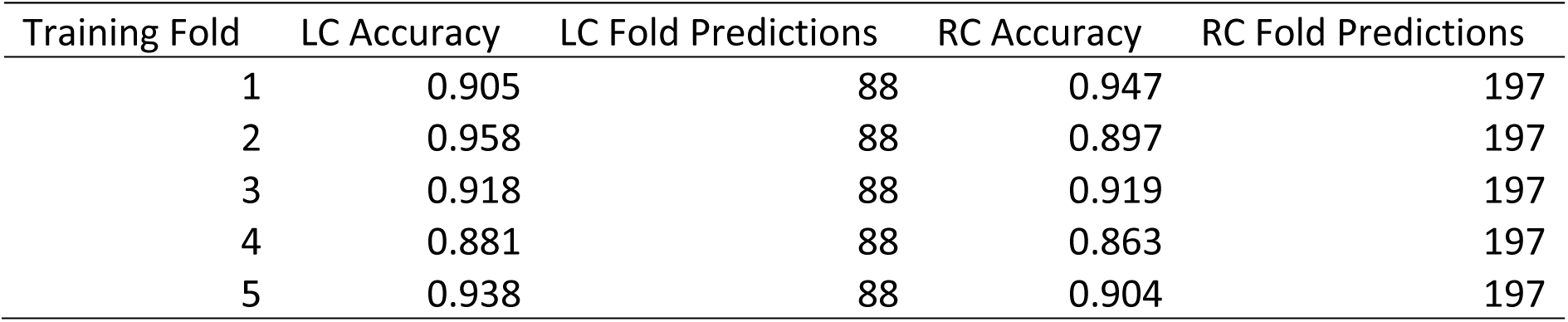
Gaussian mixture model training and cross validation for left colonic (LC) and right colon (RC) CD biopsy cell type classification.

**Table S6.** Differential expression analysis results and mean gene expression (Epithelial-Epi, Goblet-Gob, Stromal-Str, T-cell) of Infliximab resistance signature genes in left and right colonic CD biopsies (Kruskal-Wallis Test).

| Gene | Biopsy | KW p-value | Epi. Mean | Gob. Mean | Str. Mean | T-cell Mean |
| --- | --- | --- | --- | --- | --- | --- |
| MAP3K1 | Right | 0.000 | 0.144 | 0.433 | 0.677 | 0.157 |
| ACTN1 | Right | 0.079 | 0.013 | 0.017 | 0.016 | 0.066 |
| LAMA4 | Right | 0.425 | 0.128 | 0.140 | 0.092 | 0.388 |
| LAMB1 | Right | 0.000 | 0.022 | 0.017 | 0.040 | 0.388 |
| LAMB2 | Right | 0.227 | 0.007 | 0.006 | 0.020 | 0.008 |
| LAMC3 | Right | 0.174 | 0.110 | 0.067 | 0.124 | 0.083 |
| GRB2 | Right | 0.632 | 0.006 | 0.006 | 0.012 | 0.017 |
| PARVB | Right | 0.001 | 0.410 | 0.354 | 0.363 | 0.174 |
| RND3 | Right | 0.310 | 0.211 | 0.253 | 0.259 | 0.248 |
| COL4A1 | Right | 0.000 | 0.123 | 0.275 | 0.223 | 0.851 |
| ILK | Right | 0.001 | 0.104 | 0.079 | 0.036 | 0.050 |
| COL4A2 | Right | 0.195 | 0.058 | 0.045 | 0.116 | 0.116 |
| COL5A2 | Right | 0.366 | 0.013 | 0.011 | 0.004 | 0.000 |
| ITGA1 | Right | 0.000 | 0.021 | 0.028 | 0.064 | 0.157 |
| COL1A1 | Right | 0.077 | 0.004 | 0.017 | 0.016 | 0.008 |
| COL12A1 | Right | 0.163 | 0.123 | 0.096 | 0.167 | 0.091 |
| COL1A2 | Right | 0.024 | 0.055 | 0.090 | 0.072 | 0.008 |
| LAMC1 | Right | 0.011 | 0.123 | 0.112 | 0.120 | 0.017 |
| MAP3K1 | Left | 0.000 | 0.446 | 0.316 | 1.015 | 0.412 |
| ACTN1 | Left | 0.370 | 0.007 | 0.018 | 0.000 | 0.020 |
| LAMA4 | Left | 0.382 | 0.055 | 0.070 | 0.067 | 0.392 |
| LAMB1 | Left | 0.000 | 0.022 | 0.009 | 0.000 | 0.275 |
| LAMB2 | Left | 0.581 | 0.007 | 0.026 | 0.015 | 0.000 |
| LAMC3 | Left | 0.012 | 0.059 | 0.132 | 0.052 | 0.059 |
| GRB2 | Left | 0.012 | 0.004 | 0.018 | 0.000 | 0.039 |
| PARVB | Left | 0.202 | 0.531 | 0.491 | 0.418 | 0.431 |
| RND3 | Left | 0.001 | 0.498 | 0.596 | 0.478 | 0.137 |
| COL4A1 | Left | 0.164 | 0.089 | 0.158 | 0.119 | 0.196 |
| ILK | Left | 0.392 | 0.042 | 0.044 | 0.022 | 0.078 |
| COL4A2 | Left | 0.103 | 0.070 | 0.026 | 0.097 | 0.020 |
| COL5A2 | Left | 0.807 | 0.013 | 0.009 | 0.007 | 0.000 |
| ITGA1 | Left | 0.228 | 0.031 | 0.070 | 0.045 | 0.078 |
| COL1A1 | Left | 0.809 | 0.004 | 0.009 | 0.007 | 0.000 |
| COL12A1 | Left | 0.005 | 0.070 | 0.114 | 0.157 | 0.157 |
| COL1A2 | Left | 0.263 | 0.065 | 0.035 | 0.037 | 0.118 |
| LAMC1 | Left | 0.000 | 0.312 | 0.386 | 0.075 | 0.020 |

## Acknowledgments

The authors wish to thank Elizabeth A. Proctor and Alina Starchenko for their helpful comments on the manuscript.

## Author contributions

Conceptualization: DKB and DAL. Methodology: DKB and MPK. Formal Analysis: DKB. Validation: PNV, ANS, AJS, EAS, LAC, and KTW. Writing: DKB, MPK, and KSL, DAL. **Competing interests**: The authors have no competing interests to declare. **Data and materials availability**: Single-cell data is deposited in Gene Expression Omnibus (**GEO Accession Number GSEXXXXXXX).**

